# EEG-based Prediction of Cognitive Load in Intelligence Tests

**DOI:** 10.1101/539486

**Authors:** Nir Friedman, Tomer Fekete, Ya’akov (Kobi) Gal, Oren Shriki

## Abstract

Measuring and assessing the cognitive load associated with different tasks is crucial for many applications, from the design of instructional materials to monitoring the mental wellbeing of aircraft pilots. The goal of this paper is to utilize EEG to infer the cognitive workload of subjects during intelligence tests. We chose the well established advanced progressive matrices test, an ideal work-frame because it presents problems at increasing levels of difficulty, and has been rigorously validated in past experiments. We train classic machine learning models using basic EEG measures as well as measures of network connectivity and signal complexity. Our findings demonstrate that cognitive load can be well predicted using these features, even for a low number of channels. We show that by creating an individually tuned neural network for each subject, we can improve prediction compared to a general model and that such models are robust to decreasing the number of available channels as well.

## 1 INTRODUCTION

The performance of complex tasks requires the integration of various mental resources, such as task-related knowledge, working memory, attention and decision making. However, our brains have limited resources for processing and integrating information. The concept of cognitive load generally refers to the relative load on these limited resources (Sweller et al., 1998; Coyne et al., 2009).

Cognitive workload has been explored from different perspectives. Brouwer et al. (2012) refer to workload as the working memory load in an n-back task. Mills et al. (2017) use simple true-false questions for eliciting low workload and open-ended questions, which require more precise memory, for eliciting high workload. Other studies have emphasized the role of skill acquisition in modeling cognitive load (Sweller et al., 1998). Logan (1985) show that when subjects acquire a skill and learn how to perform a task in an automatic manner, their cognitive workload decreases. Thus, the cognitive load depends not only on task complexity but also on subject skill at the given task. A highly complex task performed by a non-skilled individual would result in high cognitive load, whereas a simple task performed by a skilled individual would result in low cognitive load. For example, Stevens et al. (2006) assessed subjects as they were learning to diagnose disorders of organ systems and Mak et al. (2013) focused on performance improvement in a visual-motor task. Both studies showed a decrease in cognitive load metrics, with an increase in task familiarity. In all of these studies, the difficulty of the task is seen as a proxy for its associated cognitive load. The difficulty was assessed using a variety of approaches, such as the type of questions (true-false vs. open ended), subject performance and even participant subjective ratings. We chose a domain in which problem difficulty was rigorously validated and is commonly used in the psychological literature (see below).

The focus of this paper is on quantifying cognitive workload using electroencephalography (EEG). Several studies have previously developed EEG-based measures for cognitive load. In particular, it was found that the ratio between the theta power (4-8 Hz) and the alpha power (8-12 Hz), as well as the ratio between the beta power (12-30 Hz) and the alpha power and several related combinations, provided informative indices concerning task engagement and cognitive workload (Pope et al., 1995; Mills et al., 2017; Stevens et al., 2006). Other researchers came to similar conclusions, that the relation between different spectral features can help predict cognitive load from EEG (Gerě and JauĆcvec, 1999; McDonald and Soussou, 2011; Conrad and Bliemel, 2016). This study aimed to further expand these studies and develop more accurate EEG-based measures of cognitive load.

We focused on recording EEG during performance of a well-known psychological assessment tool, the advanced progressive matrices test (Raven, 2000), which is commonly used to measure general intelligence. The test is composed of different problems that involve the manipulation of shapes. Each problem differs in difficulty level, which was independently validated across a large number of subjects (Forbes, 1964; Arthur Jr et al., 1999). Problems are presented to subjects at increasing level of difficulty. The difficulty of each problem is validated across a large number of subjects (Forbes, 1964; Arthur Jr et al., 1999).

Here, we adopt this as the operational definition of cognitive load. We demonstrate that it can be predicted from the subject’s EEG readings. Specifically, we use different EEG measures as input to machine-learning algorithms and train them to predict problem difficulty.

Previous studies of EEG-based measures of cognitive load were limited by several aspects. In particular, they relied mostly on spectral features and produced a simple discrete measure of either low or high load. In contrast, this paper models cognitive load in a continuous manner. In addition, we go beyond basic spectral features and examine how measures of network connectivity and signal complexity affect the prediction of cognitive load. To measure network connectivity, we used complex network analysis (CNA) which provides measures to examine functional connectivity in the brain (Bullmore and Sporns, 2009; Fekete et al., 2014). Features of neural complexity are often computed using measures of entropy, reflecting the proportion of ordered patterns that can be detected in a signal (Bullmore et al., 2009). To measure neural complexity, we focused on Lempel-Ziv (Tononi and Edelman, 1998) complexity, Multi Scale Entropy (MSE) (Abásolo et al., 2006) and Detrended Fluctuation Analysis (DFA) (Rubin et al., 2013).

The results of this paper demonstrate the applicability of using EEG and machine learning for quantifying cognitive load in well validated problem-solving tasks. In particular, as EEG and other measures of brain activity become more pervasive, quantitative cognitive load measures could be used to facilitate the design of domains involving real-time problem-solving, such as e-learning, psychometric exams, military training, and more (Mills et al., 2017; Ikehara and Crosby, 2005).

## 2 METHODS

We recorded EEG from subjects while they solved the Advanced Progressive Matrices set II (Raven test). The 36 problems in the test were presented in increasing level of difficulty. The raw EEG data were then passed through an artifact removal pipeline (see details below), before extracting EEG-based measures of spectral activity, neural complexity and network connectivity. These measures served as input to machine learning algorithms, which were trained to predict problem difficulty.

### 2.1 Participants

Fifty-two subjects (26 female and 26 male; age range 21 - 28, *Mean* = 24.55 years, *SD* = 1.76 years) participated voluntarily in the experiment, provided written informed consent and received compensation for participating. The experiment was approved by the Ben-Gurion University ethics committee. All subjects reported that they are right-handed, have normal or corrected vision, and that they have never completed any sort of intelligence test in the past. Four participants required less than ten minutes to solve the entire test, and/or correctly answered 16 problems or less, and were excluded from further analysis. In addition, one subject’s recording was compromised dye to technical difficulties, and was excluded as well.

### 2.2 Experimental paradigm

Subjects performed the Raven’s APM Set II problems (36 items in increasing difficulty level), and instructions were delivered before the test started (see Figure 1 for an example problem). The test was run with no time limit, with all the key requirements and administration instructions carefully following the manual (Raven et al., 1998). Subjects sat in a comfortable chair, facing a computer screen 60 cm away. The test was conducted by displaying the problems on the computer screen (23”, 1920 × 1080 resolution, with a 2.3° visual angle between each answer’s corners), where the subjects were required to press a keyboard key (with their right hand) in accordance with their chosen answer number. The experiment was programmed in MATLAB (http://www.mathworks.com, version 2015), using the Psychophysics Toolbox extensions (Brainard and Vision, 1997; Pelli, 1997; Kleiner et al., 2007). Each trial lasted from the presentation of the corresponding problem until the subject response, and thus trial duration was variable.

**Figure 1.**
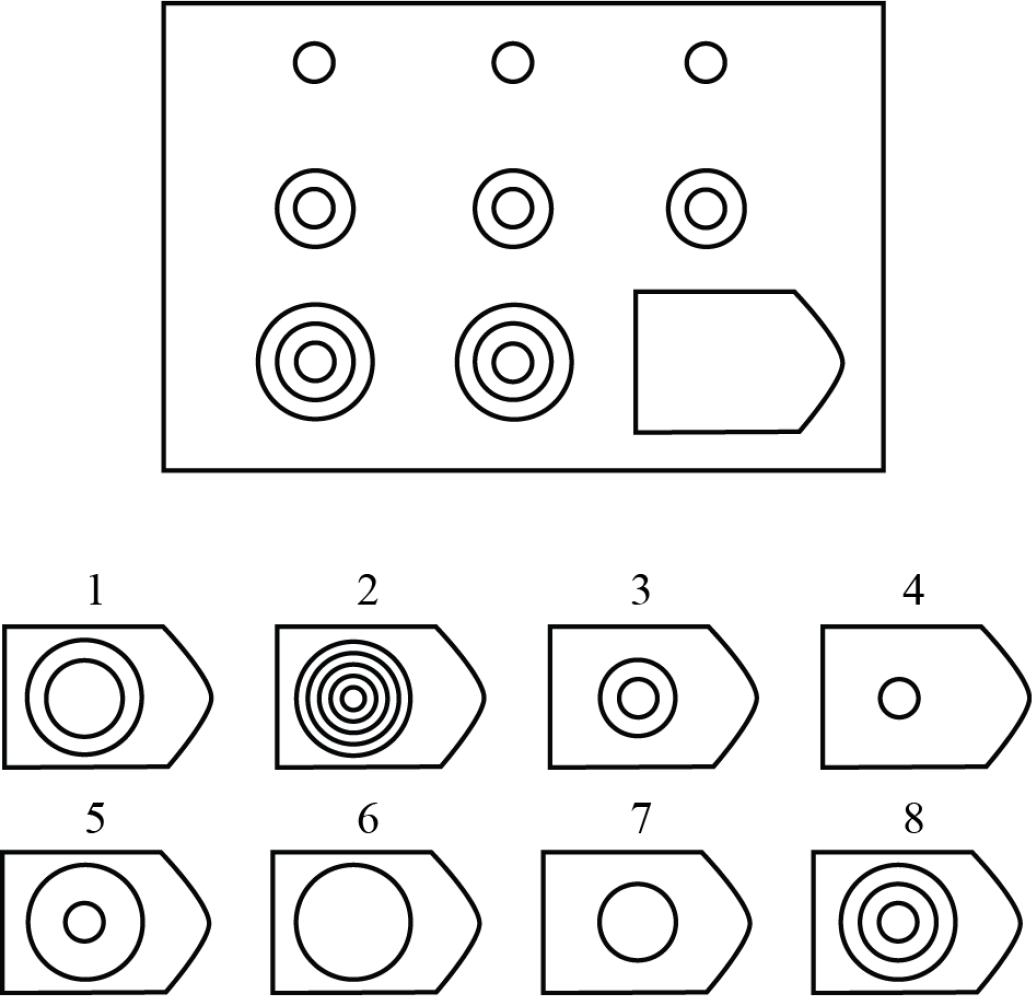
Illustration of Raven’s set II Example Problem. The subject is asked to choose the missing shape from the 8 possible options. The correct answer here is option 8.

EEG was recorded through the whole session using the g.Tec HIamp system (g.Tec, Austria) with 64 gel-based electrodes (AgCl electrolyte gel). Electrodes were positioned according to the standard 10/20 system with linked ears reference. An impedance test and adjustment was carried out at the beginning of the session, and impedances of all electrodes were kept below 5 kΩ. The signal was sampled at 256 Hz with a high-pass filter of 1 Hz. The data were recorded using Matlab Simulink g.Tec plug-ins.

### 2.3 Feature extraction

Data were analyzed using a combination of the EEGLAB Matlab plug-in (Delorme and Makeig, 2004) routines and custom code. Data were first high-pass filtered (cut-off 1Hz), then a customized adaptive filter was applied to suppress line-noise. This followed by Artifact Subspace Reconstruction (Mullen et al., 2015), re-referencing to the mean, and low-pass filtering (cutoff 60Hz). Next, Infomax ICA was carried out. The resulting ICs were evaluated automatically for artifacts, by combining spatial, spectral and temporal analysis of ICs. ICs identified as containing ocular, muscular or cardiac artifacts were removed from data.

Various features were extracted from the EEG data:

- **Power spectrum metrics (PS) -** The power in 5 frequency bands (delta [1-4 Hz], theta [4-8 Hz], alpha [8-12 Hz], beta [12-30 Hz] and gamma [30-50 Hz]) was calculated for each channel across the whole trial duration. This resulted in 310 features (62 channels × 5 bands) for each trial.
- **Neural complexity metrics -** We focused on three measures of complexity, specifically, Lempel-Ziv complexity (LZC) (Zhang et al., 2001), Multi Scale Entropy (MSE) (Abásolo et al., 2006) and Detrended Fluctuation Analysis (DFA) (Peng et al., 1995; Rubin et al., 2013). The LZC measure was computed as the mean of the measure across all channels, resulting in a single feature for each trial. In comparison, the MSE and DFA measures were first computed for each individual channel, and for DFA we also computed the metric for each frequency band (as described above). We then computed the mean, variance, maximum, minimum, 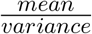 and 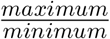, resulting in 6 features for the MSE, and 36 features for DFA (6 measures × (5 bands + 1 broadband)), resulting in 43 complexity features for each trial. Because these metrics are affected by trial duration, we calculated them for the last 2500 samples (≈10 seconds) of each trial.
- **Connectivity metrics -** These features are based on a graph reflecting the connectivity of the underlying network. The graph includes 62 vertices (channels); edges in the graph represent correlations between channels (there are no self edges). There are two approaches regarding the weight of each edge. One is to take the absolute value of the correlation as the weight of each edge. Another is to give the same weight to all edges that were kept after the thresholding process described below. We kept only the top x% (we tried several thresholds) of the edges with the highest values, for example 5%, meant that we were left with 190 edges out of the 62^2^ (minus the 62 self edges). The graph was used to extract graph-theoretical features such as longest/shortest distance between nodes, small-worldness etc (Bullmore and Sporns, 2009). We ultimately used the mean and standard deviation of the small-worldness measure and its components, across the different thresholds.
- **Basic -** Simple demographic features of subject’s age and sex were used. In addition, the time it took to answer each problem was used as a feature. These features were added to all the above feature groups in the prediction phase.

## 3 RESULTS

After removal of subjects who did not meet the inclusion criteria (see Methods), we were left with 47 subjects for the analysis (24 female and 23 male; age range 21 - 28, *Mean* = 24.55 years, *SD* = 1.79 years). Our goal was to estimate the cognitive workload of subjects as they were trying to solve each problem during the test.

To this end, we assumed that the difficulty level increased with every problem, as validated in previous studies (Forbes, 1964; Arthur Jr et al., 1999). Figure 2 shows the rate of incorrect responses over all problems in our data, reflecting the established relationship between problem number and difficulty level. Interestingly, problems 24 and 29 deviated significantly from the trend (more than 3 standard deviations). For this reason, both problems were also excluded from our analysis. In addition, we only considered trials where subjects answered correctly. This is based on the premise that in most cases if the subject answered incorrectly, then he or she might have not reached the appropriate load level. After excluding the subjects (180 trials) and specific problems (94 trials), as stated above, and the incorrect trials (366 trials), we were left with 1232 trials.

**Figure 2.**
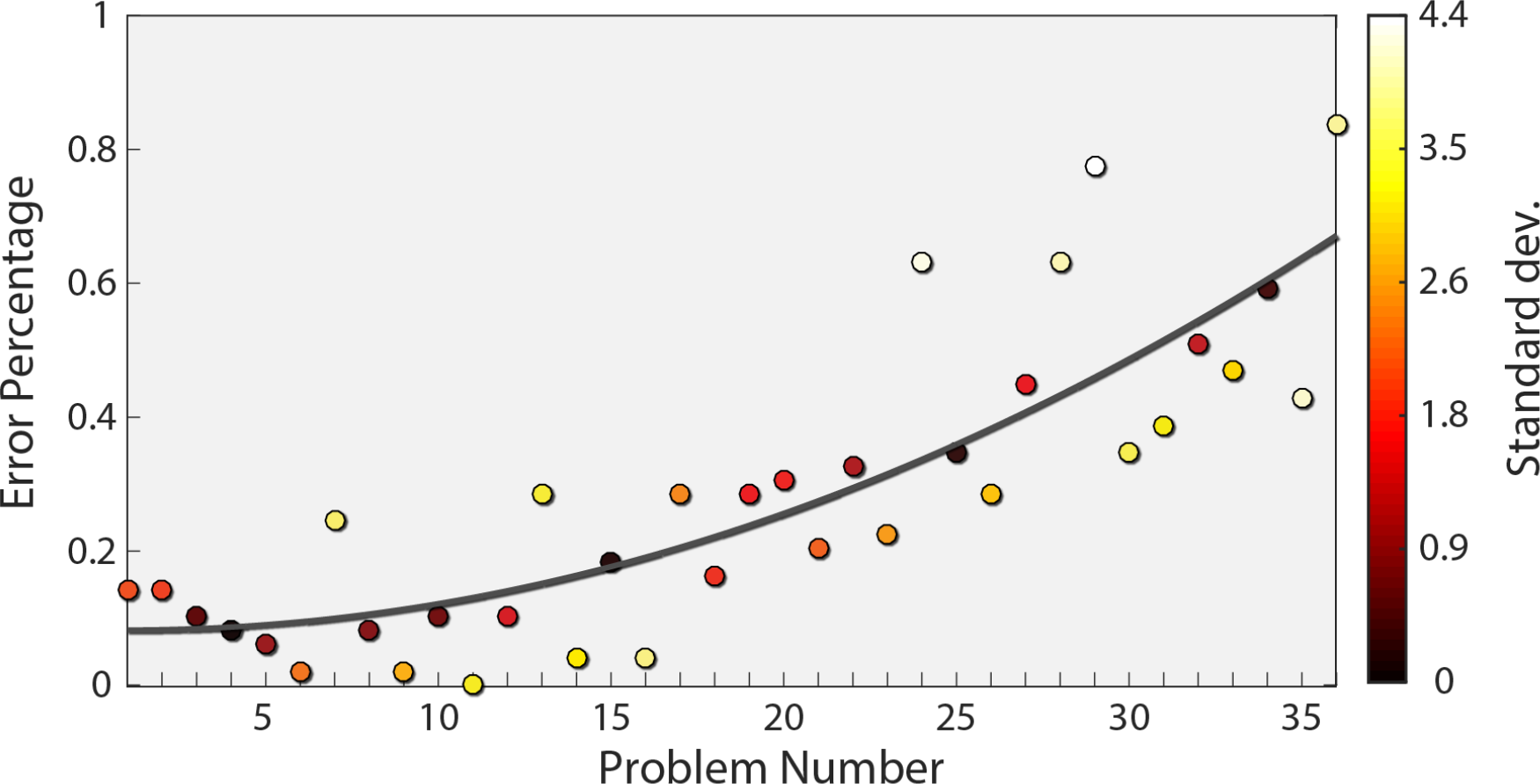
Subject Error rate as a function of problem number. The mean error rate across subjects is plotted for each problem (circles) together with a quadratic fit (red curve). The number near each point indicates the number of standard deviations from the fit.

For each of the 1232 correct trials, we computed different features (as detailed in the Methods) and assigned it with the corresponding difficulty level (a number between 1 and 36) as the target value. Several types of machine learning algorithms were tested in order to predict cognitive load - ‘Random Forest’ (RF) from the sklearn python package (Buitinck et al., 2013) and ‘XGBoost’ (XGB) and it’s corresponding python package (Chen and Guestrin, 2016). They were chosen because of the virtues of an ensemble learning algorithm, along with their usual good fit with temporal data. Additionally, we applied an artificial neural network (ANN), using the keras python package (Chollet et al., 2015). Lastly, we used simple Linear Regression (LR), also from the sklearn python package, as a baseline for comparison. The hyper-parameters of these models were found using a grid search. The best performance was exhibited by the XGBoost classifier with a step size of 0.05. For the optimal feature group, the number of boosting rounds was 300. All other parameters were run with the default settings. All results shown were cross-validated by dividing the data randomly to training and validation sets (80% of the data were used for training, 20% of the data were used for validation) and repeating the process 10-20 times.

At first, we compared the different feature types in the prediction process with the different classifiers. Table 1 summarizes the *r*^2^ results for this analysis (all results marked with an * were significant). The best results were obtained using XGB for all feature types. XGB provides a good trade-off between model complexity and the number of samples required to reach robustness. Even though ANN can capture very complex relationships, they require a large training set. On the other hand, LR and RF do not require significant amounts of training data, but their model complexity is significantly more constrained than XGB.

**Table 1.** The table shows the Pearson correlation (*r*^2^) of each Feature group - Model Pair.

|  | **LR** | **RF** | **XGB** | **ANN** |
| --- | --- | --- | --- | --- |
| **PS** | 0.007 | 0.383* | 0.655* | 0.346* |
| **Complexity** | 0.323* | 0.055 | 0.508* | 0.286* |
| **Connectivity** | 0.335* | 0.186 | 0.5* | 0.267* |
| **PS & Complexity** | 0 | 0.322* | 0.641* | 0.186* |
| **PS & Connectivity** | 0.07 | 0.44* | 0.67* | 0.32* |
| **Complexity & Connectivity** | 0.339* | 0.122 | 0.519* | 0.331* |
| **All Features** | 0.05 | 0.358* | 0.628* | 0.297* |

Next, we compared the utility of each of the three different feature types. PS and connectivity features obtained the highest score, and adding the complexity features to either of the two, did not contribute significantly to the prediction. This suggests that complexity features do not add any further information beyond spectral features and connectivity features. To test whether this was not due to high model complexity resulting in over-fitting, we conducted a feature selection process. We found that even after reducing the number of features, no combination of complexity, connectivity and PS features yielded better results than using only the PS and connectivity features together with the basic features.

Figure 3 shows a scatter plot of the best model’s prediction together with the true label of each instance in the test set. The Pearson correlation of the best model is *r*^2^ = 0.67 (*p* < 0.01). The model was trained on the problem serial number, which should, in principle, produce a linear relationship. However, as evident in Figure2, the relationship between problem number and error rate is slightly non-linear. This suggests that the relationship between problem number and the EEG measure could also be non-linear. We therefore also computed the Spearman correlation, which relates to a general monotonic relationship rather than a linear one, and obtained a value of 0.81 (*p* < 0.01). One of features used by the algorithm was the duration of each segment, namely the time it took the subject to answer. We also examined the performance with only this feature and found a of *r*^2^ = 0.23 (*p* < 0.01) and a Spearman correlation of 0.41 (*p* < 0.01).

**Figure 3.**
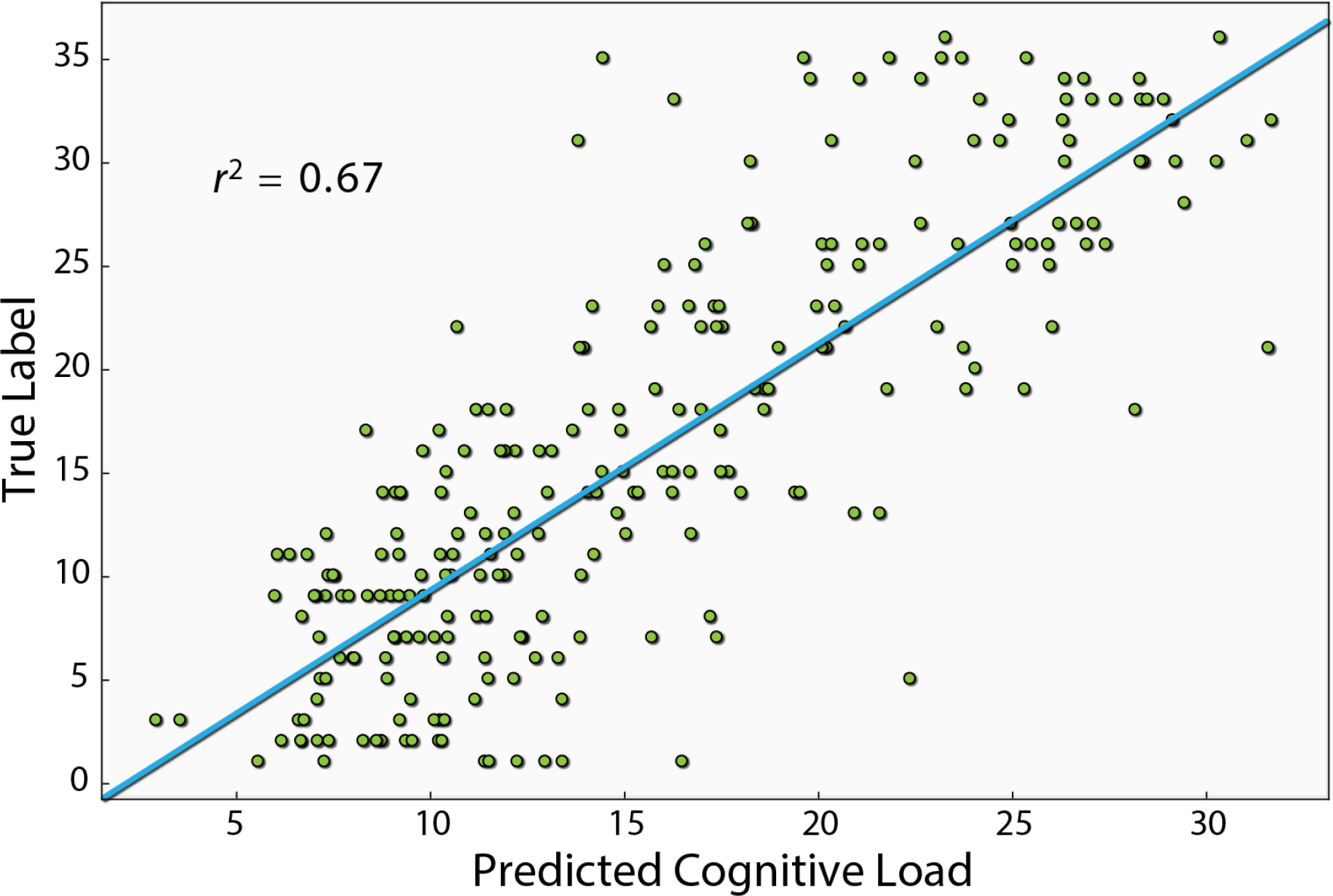
This figure shows the Pearson correlation between the XGBoost model’s prediction and the true label of each instance. The model shown here uses the PS, connectivity and basic features, which is the one that produced the best prediction.

### 3.1 Effect of number of electrodes

From an applicative point of view, the number of electrodes affects both the cost and the complexity of using EEG. We therefore examined the extent to which reducing the number of electrodes affects the prediction quality. To this end, we conducted a two step analysis. Firstly, we ran 1,000 simulations, where in each, ten electrodes were chosen randomly out of a total of sixty-two. For each electrode combination, only the relevant PS features were used (five per channel, in addition to the basic features) to generate a workload prediction using the XGB algorithm. We then sorted the electrodes based on the percentage of simulations each electrode was involved in that yielded a score above a specified threshold, out of the entire simulations it participated in. The top thirty electrodes were chosen in descending order, and were taken for the second step, where the effect of the number of best electrodes on the *r*^2^ was examined. As seen in Figure 4, a relatively high *r*^2^ of 0.7 (*p* < 0.01) can be obtained using only 12 electrodes (and in fact over 95 percent of peak performance for only 8). Additionally, using the same features of the 12 electrodes, the model produces a Spearman correlation of 0.82 (*p* < 0.05).

**Figure 4.**
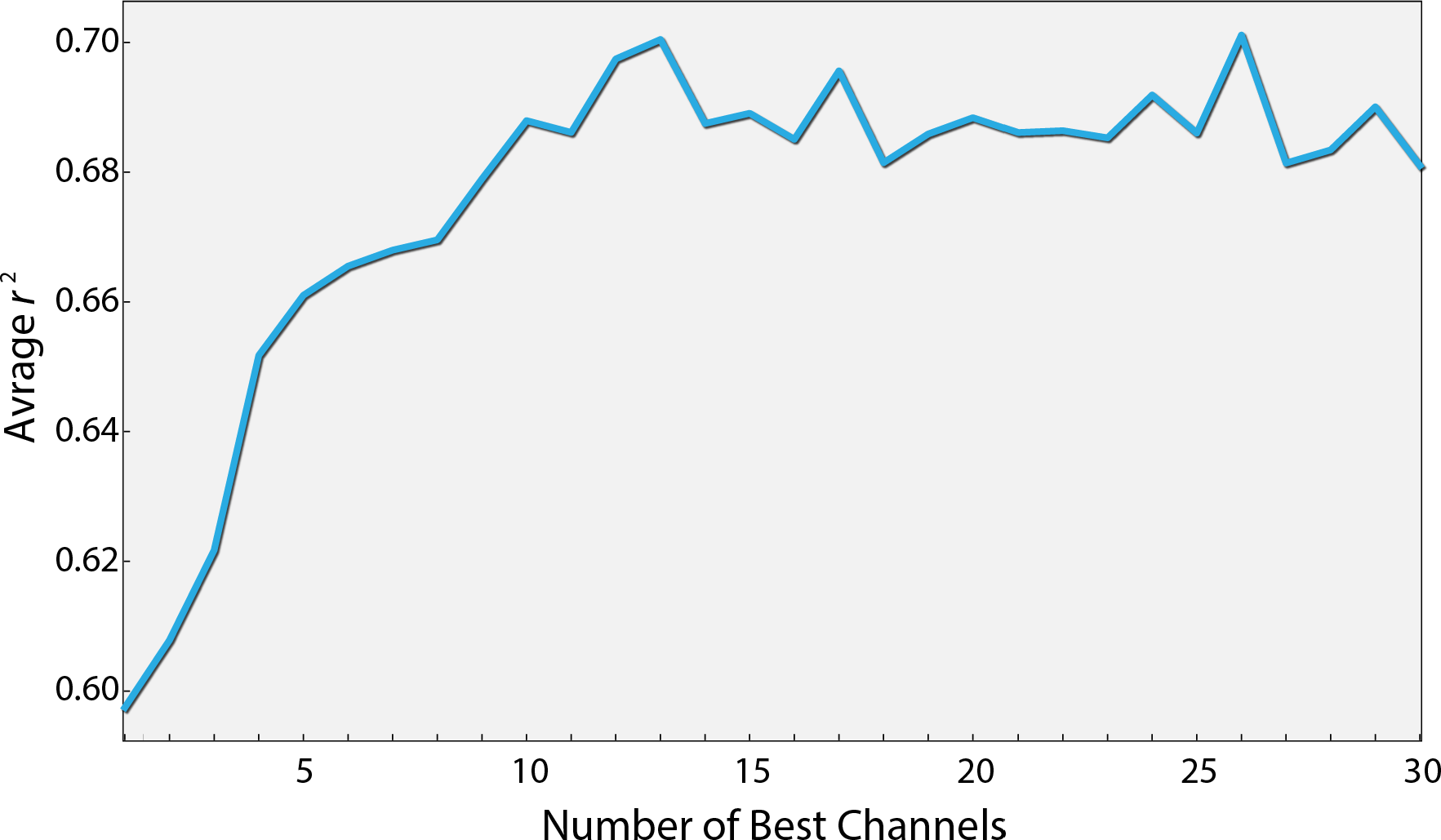
Performance as a function of the number of best channels. Channels were ordered according to their contribution to the prediction quality (see text for details). The curve depicts the prediction quality (*r*^2^) for the XGBoost algorithm as a function of the number of best channels taken into account.

### 3.2 Effect of discretizing the workload

In our analysis, the target variable (difficulty of each problem) had 34 different values. We analyzed the influence of reducing the number of levels of the target variable. We used different sized bins, to reduce the number of different values to 6, 9, 18, 34. For example, to obtain 6 levels, values were binned to [1-6], [7-12], [13-18], [19-24], [25-30] and [31-36]. As evident in Figure 5, prediction quality generally decreased with the number of levels. This is not surprising, because the prediction task becomes more complex with the number of levels. In addition, we show that using only the best 12 electrodes found earlier to compute the connectivity features (combined with the PS features of those electrodes), we obtain *r*^2^ = 0.713 (*p* < 0.05) for 6 levels, which is the best prediction quality we obtained.

**Figure 5.**
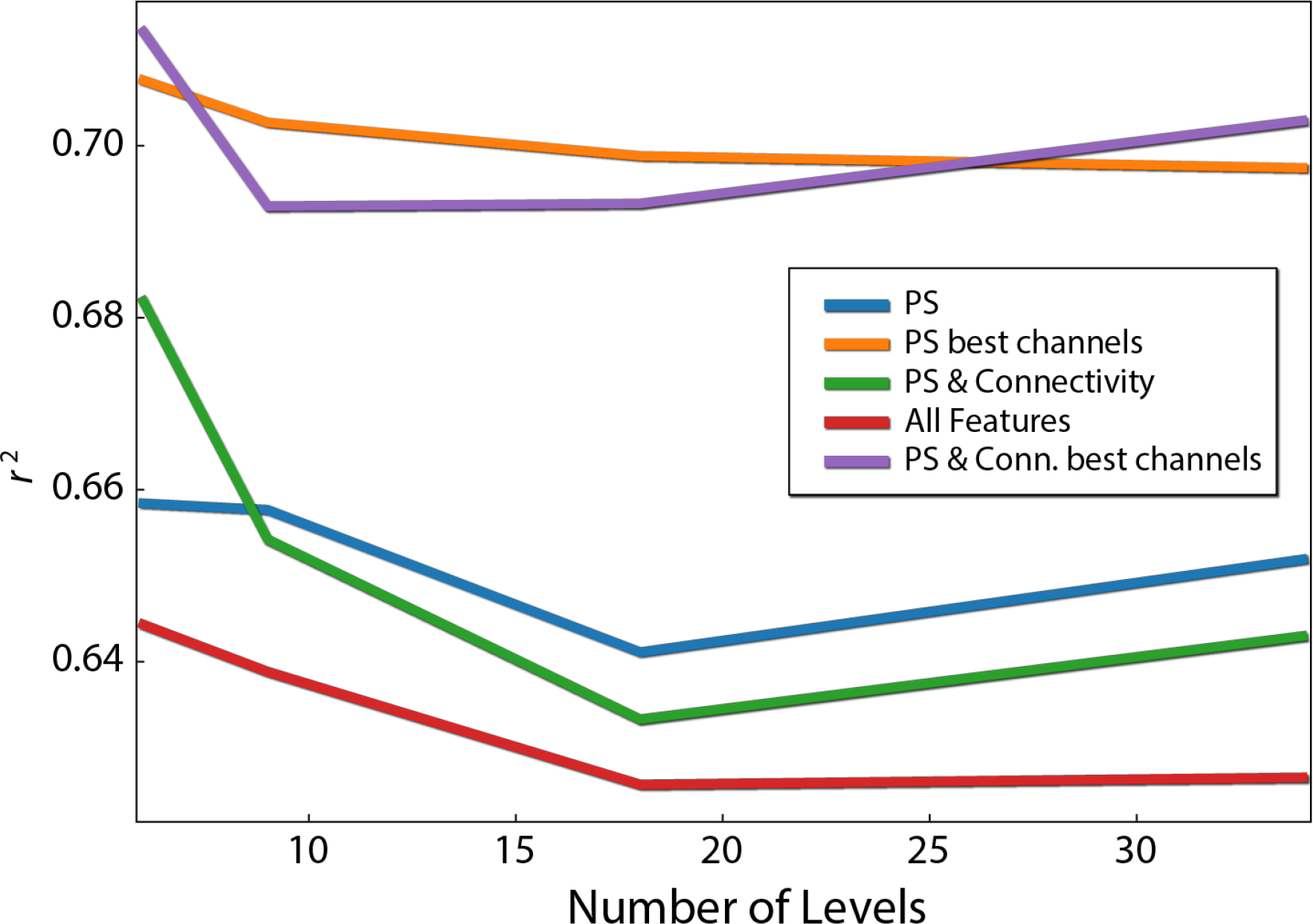
Difficulty level discretization effect on prediction quality (*r*^2^). Each line corresponds to different feature types. PS red are the PS features of the 12 best channels.

### Individualized prediction using neural networks

Lastly, because different individuals might experience different levels of cognitive load for the same problem, we wanted to assess the influence of individualizing the prediction model. To this end, we first built a three layer artificial neural network (ANN), trained with data from all subjects using the PS and connectivity features of the 12 best electrodes. We then fixed the parameters of the first and second layers, and for each subject continued to train the weights of the output layer (Figure 6). We conducted a paired t-test (Figure 7), by calculating the mean correlation with the correct answer over several folds using the general model (*M* = 0.39, *SD* = 0.06) and after tuning (*M* = 0.43, *SD* = 0.06), which yielded a significant difference (*t* = −4.75, *p* = 0.001) in favor of the individualized network models.

**Figure 6.**
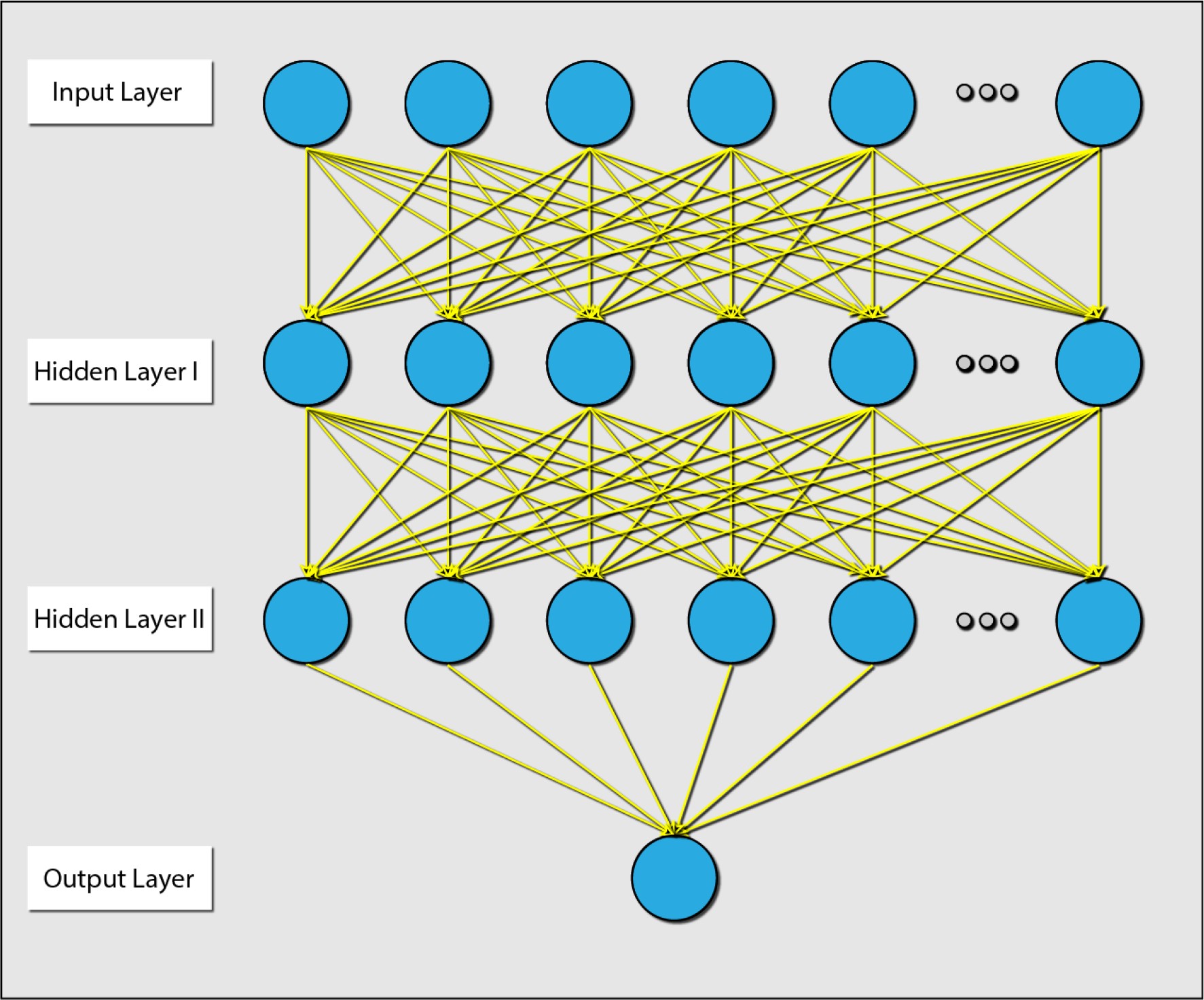
Diagram explaining the architecture of the ANN that was used. There were 2 hidden layers, and all layers were dense (e.g. all connections were present). The parameters between the input layer and hidden layer 1, and the parameters between hidden layer 1 and hidden layer 2 were held during the individualization phase.

**Figure 7.**
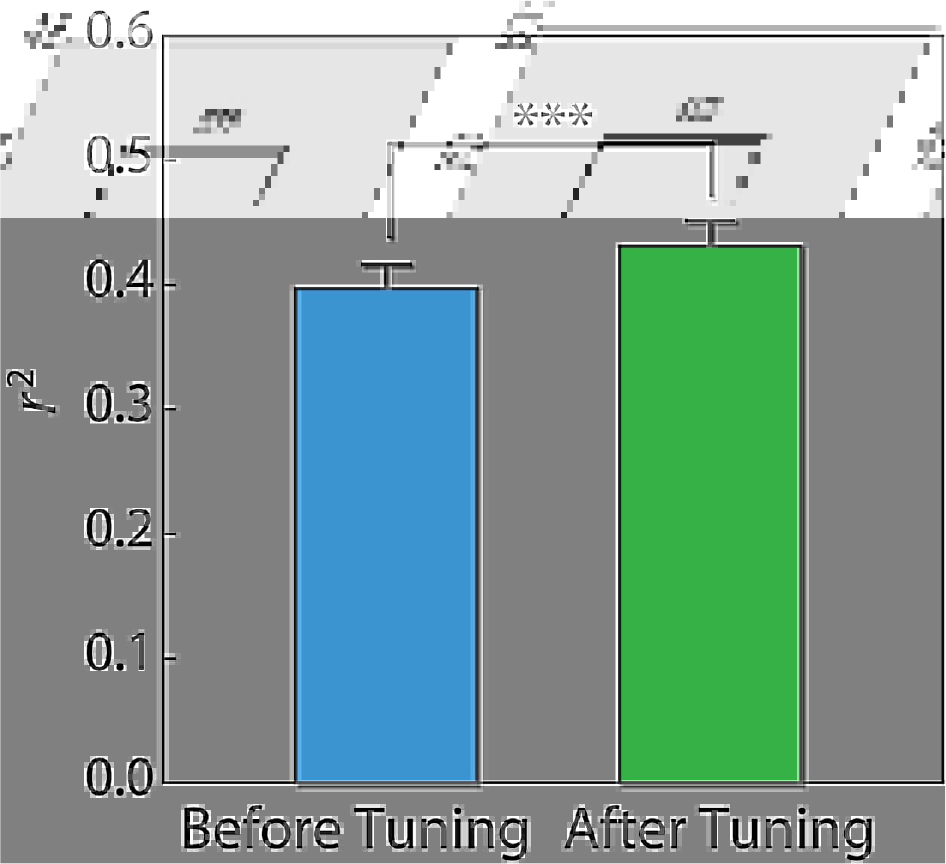
Difference of *r*^2^ score using an ANN before and after individualizing the last layer for each subject. Error bars reflect standard error of the mean. Paired t-test results are shown.

## 4 DISCUSSION

We recorded EEG from subjects while solving the advanced progressive matrices test (Raven’s matrices test), and used EEG features and machine learning to predict problem difficulty, which we interpreted as cognitive workload. Problem difficulty was ordered on a scale from 1 to 36 (Forbes, 1964; Arthur Jr et al., 1999) and was treated here as a continuous value. Our results show that even when considering cognitive load in a continuous manner, a reasonable prediction accuracy can be obtained using EEG measures. This could be very useful for many applications in which there is a wide range of cognitive workload levels. These findings extend those of previous studies which used a small number (2-3) of discrete levels of cognitive workload (Gerě and JauĆcvec, 1999; McDonald and Soussou, 2011; Conrad and Bliemel, 2016). Indeed, we found that reducing the number of difficulty levels, improves the results significantly.

We examined several machine learning algorithms and found that XGBoost outperformed all other algorithms with all three feature groups. XGBoost was more accurate than the simpler models of linear regression and Random Forest. The lower scores of the ANN are probably due to the fact they typically require a much larger training dataset than we had at our disposal (Chen and Guestrin, 2016). Furthermore, even though the ANN scored lower than XGBoost, we showed that prediction quality can be improved by tuning the last layer of the ANN to each individual. With a larger dataset, the personalized ANN could potentially attain better prediction than XGBoost.

As part of our analysis, we checked the impact of additional EEG measures, specifically metrics of connectivity and metrics of neural complexity. Our results suggest that connectivity measures do add information regarding cognitive load, beyond the simple spectral features. On the other hand, it seems that complexity features, while holding information regarding cognitive load, do not explain anything more than connectivity and PS features.

Lastly, we found that prediction quality did not deteriorate, and even improved, when using a limited number of channels (~12), which is important for practical applications. This is most probably due to better generalization, resulting from a less complex model, as opposed to one utilizing all channels.

We chose to utilize the advanced progressive matrices test in this study because of the high validity of its operationalization of difficulty levels. However, to extend our findings further towards applicability, future studies should examine the utility of our EEG based metrics for cognitive load in real-life settings such as control tower operator performance as aerial traffic ebbs and flows. Since our results indicate the feasibility of employing an array comprising as little as eight electrodes, potentially such studies could be carried out in parallel using portable dry EEG systems. The added benefit would be the feasibility of amassing the expansive datasets necessary for utilizing elaborate neural network models, which in this scenario are expected to improve predictive ability. In addition, it would be useful to identify EEG markers for different dimensions of cognitive workload. Such markers would pave the way for optimizing and personalizing learning processes from e-learning to military training (Mills et al., 2017; Ikehara and Crosby, 2005).

## AUTHOR CONTRIBUTIONS

Experiment planning and design- N.F., K.G., O.S.

Data acquisition- N.F.

Data analysis- N.F., T.F.

Writing the manuscript- N.F., T.F., K.G., O.S.

## ACKNOWLEDGMENTS

The authors thank Dr. Nir Getter for helpful discussions on the manuscript and Ms. Koral Regev for valuable help in running the experiments.

